# Postnatal Abrogation of VEGFR2 Blocks Terminal Cap2 Differentiation by Preventing the Developmental Progression from a Capillary Intermediate Cell State

**DOI:** 10.1101/2025.07.23.666389

**Authors:** Daoqin Zhang, Carsten Knutsen, David J. Stroud, Xibing Che, David N. Cornfield, Fabio Zanini, Cristina M. Alvira

## Abstract

After birth, the alveolar capillary network expands to increase gas exchange surface area and endothelial-derived signals promote alveolarization. Lung capillaries are comprised of two distinct subsets, one with proliferative potential to facilitate growth and repair (Cap1), and the other serving a specialized role in gas exchange (Cap2). However, the molecular mechanisms directing capillary speciation, developmental plasticity, and fate transitions during development and repair are not well understood. Here, we show that Cap2 are absent in late embryonic life but rapidly appear and expand immediately after birth. We show that Cap1 progenitors first transition to a novel, intermediate cell state (Cap^INT^), characterized by co-expression of Cap1 and Cap2 markers, and heightened proliferation. Cap^INT^ are present in both the developing mouse and human lung. Hyperoxia, an experimental model of bronchopulmonary dysplasia (BPD), a chronic lung disease marked by impaired alveolarization, increases Cap^INT^ abundance and persistence and expands Cap2 EC. Cap^INT^ EC are also increased in human infants dying with active BPD. Using genetic lineage tracing, single cell transcriptomics, ATAC-sequencing and a mouse model that permits inducible deletion of VEGFR2 in Cap^INT^ and Cap2 EC, we show that postnatal abrogation of VEGFR2 markedly increases Cap^INT^ EC abundance, blocks Cap2 terminal differentiation, impairs alveolarization, and activates alveolar fibroblasts. Finally, we identify ERG as a putative VEGFR2-downstream mechanism that promotes Cap^INT^ to Cap2 differentiation. Taken together, our data show that Cap1-Cap2 differentiation is a two-step process that only requires VEGFR2 for the second step. Elucidation of the physiologic and molecular pathways that control the initial transition of Cap1 to Cap^INT^ EC has the potential to reveal new therapeutic targets for lung diseases that disrupt the alveolar capillary formation and integrity.

## Introduction

Significant lung growth occurs after birth. Septation of distal lung saccules into millions of alveoli and rapid expansion of lung capillaries markedly increases gas exchange surface area. Pulmonary endothelial cells (EC) play a central role in promoting alveolar growth^1–3^ and lung repair after injury^4–6^, through the production of ‘angiocrine’ factors^7^. In the adult lung, injury induces EC to express growth factors that promote alveolar epithelial cell proliferation^8,9^, a process that requires signaling through the VEGFR2 receptor^9,10^, highlighting an essential role for VEGFR2 in the adaptive angiocrine response. In contrast to acute injury, chronic injury induces maladaptive angiocrine signals that activate fibroblasts and create a microenvironment favoring tissue fibrosis^11,12^.

Recent studies have shown that the alveolar capillary network is comprised of two distinct subtypes of subtype. Capillary 1 EC (Cap1, also called gCap) are smaller, express plasmalemma vesicular associated protein (*Plvap*) and KIT proto-oncogene (*Kit*), and encircle the alveolar nests. In contrast, Cap2 EC (also called aerocytes or aCap) are significantly larger, express numerous distinguishing markers (e.g. *Car4*, *Tbx2*, *Sirpa*), and extend multiple projections to form the complex vascular network covering the alveoli^13^. Interestingly, although Cap1 appear to be more proliferative than Cap2, Cap2 EC express higher levels of VEGFR2 (*Kdr*) and require alveolar epithelial type 1-derived *Vegfa* for speciation during development. However, whether VEGFR2 signaling is required postnatally, and the downstream mechanisms by which VEGFR2 promotes Cap2 differentiation remain unknown. Further, the adaptive angiocrine signals promoting postnatal alveolar formation remain poorly defined. Despite the known importance of ECs, the distinct functions of specific capillary EC subtypes in alveolar formation, their varied responses to injury, and the mechanisms governing their specialization, plasticity, and self-renewal during alveolarization remain largely unknown.

In this study, we investigated the molecular mechanisms promoting Cap2 differentiation and expansion during development and in response to injury. We show that transcriptionally distinct Cap2 EC only appear after birth, not during late embryonic development as previously described. Cap2 EC subsequently expand across early alveolarization, and hyperoxia further expands the Cap2 population. Developing a robust transcriptomic data set enriched with EC from the late saccular lung, we identified an intermediate capillary EC cell state, with Cap^INT^ sharing distinguishing markers of both Cap1 and Cap2 EC and exhibiting heightened proliferation. Given prior work identifying VEGF signaling as essential for Cap2 speciation, we developed a mouse model to permit inducible deletion of VEGFR2 from Tbx2+ EC after birth. Postnatal deletion of VEGFR2 in Tbx2⁺ EC: (i) inhibited hyperoxia-induced expansion of the Cap2 population; (ii) blocked Cap2 terminal differentiation; (iii) increased Cap^INT^ abundance; and (iv) induced activation of a fibrotic gene signature in alveolar fibroblasts. Using ATAC-sequencing to identify transcriptional regulators driving Cap2 differentiation, we found that hyperoxia selectively increased chromatin accessibility of DNA enriched in ERG-binding motifs in Cap2 EC but not in other EC, and that ERG expression was markedly decreased in hyperoxia-exposed mice subjected to Tbx2-mediated VEGFR2 deletion. Collectively, these findings show that postnatal expansion of Cap2 during development and disease requires transition to an intermediate cell state. This initial step does not require VEGFR2, however, subsequent VEGFR2-mediated activation of ERG appears essential for Cap2 terminal differentiation and the maintenance of a homeostatic alveolar capillary niche.

## Results

### CAP2 abundance markedly increases after birth and is further expanded by hyperoxia

Intense focus has centered upon the highly specialized Cap2^14^ in terms of transcriptional markers, morphology, and location^13^. However, numerous gaps in knowledge remain regarding the mechanisms that drive both initial speciation and terminal differentiation, and the specific role of Cap2 in informing the alveolar niche during development. Cap1 and Cap2 each express several distinct marker genes^15^ allowing discrimination of these two subsets. For example, *Car4* expression is significantly higher in Cap2 compared to Cap1^14,16^. In contrast, *Kit* is highly expressed by Cap1 in addition to embryonic microvascular EC, proliferative EC and some macrovascular EC, but minimally expressed in Cap2. *Tbx2*, a gene broadly expressed in mesenchymal cells, is exclusively expressed by Cap2 within the endothelial lineage (Fig. 1A).

**Figure 1:**
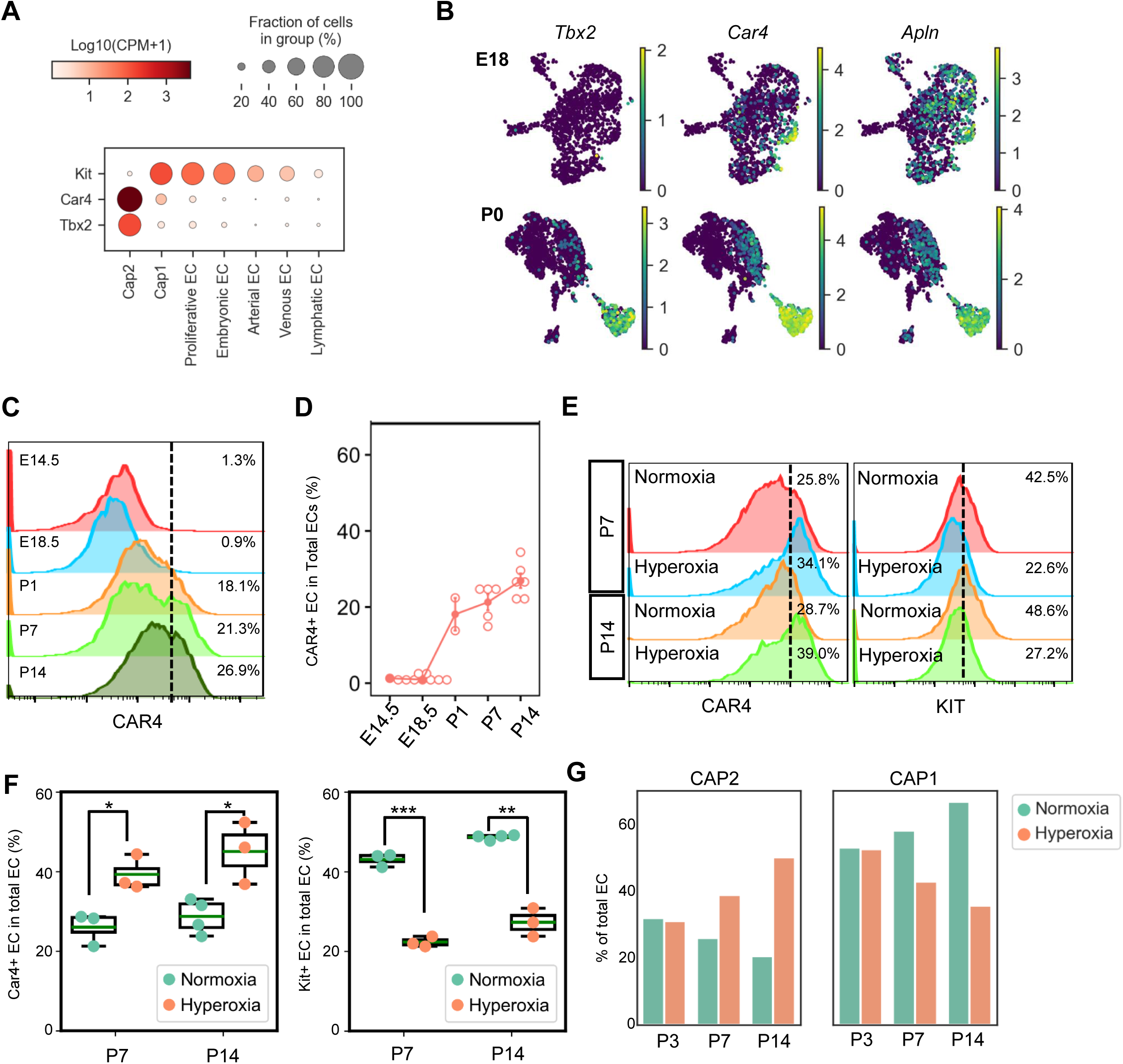
CAP2 abundance markedly increases after birth and is further expanded by hyperoxia. **(A)** Dotplot showing expression of *Kit*, *Car4* and *Tbx2* in lung EC from mice across development (P1-P21)^15^. **(B)** UMAP plots of EC from the mouse lung at E18 (top row) and P0 (bottom row) showing levels of *Tbx2*, *Car4*, and *Apln* expression from data from Negretti et al. Scale bar showing gene expression low to high (purple to yellow) in log scale counts per ten thousand **(C)** Representative FACS plot and **(D)** quantification determining the percentage of CAR4^HIGH^ EC vs. total EC at various developmental timepoints. **(E)** Quantification of FACS data to detect Car4+ Cap2 and Kit+ Cap1 in P7 and P14 neonatal mice exposed to normoxia and hyperoxia. Data are presented as mean ± SEM. Statistical significance was assessed using unpaired two-tailed Student’s *t*-test. P-values for the comparison of Car4⁺ EC percentages between normoxia and hyperoxia at P7 and P14 are 0.02 and 0.049. *P*-values for Kit⁺ EC percentages at P7 and P14 are 0.0001 and 0.008, with n=3-4 mice/group. **(G)** Re-analysis of single cell data from Hurskainen et al. quantifying the percentage of EC annotated as either Cap2 or Cap1 vs. total EC.

The earliest development timepoint corresponding to Cap2 speciation remains controversial. To clarify the emergence of Cap2 in the developing mouse lung, we re-analyzed ECs from a robust single cell dataset that includes multiple embryonic and early postnatal developmental timepoints^17^. Although there was a group of EC that expressed variable levels of *Car4* just before birth at E18, other canonical Cap2 markers (e.g. *Apln)* were not restricted to these cells, and microvascular EC at E18 completely lacked *Tbx2* expression (Fig. 1B). In stark contrast, immediately after birth (P0), a separate, transcriptionally distinct Cap2 cluster emerged that highly expressed all three Cap2 marker genes (*Tbx2*, *Car4* and *Apln*).

We validated this finding using FACS to quantify the number of CAR4^High^ EC at multiple timepoints across late embryonic and early postnatal development (Fig. 1C-D). Consistent with the transcriptomic data, at late embryonic timepoints, CAR4^High^ EC were very infrequent. By P1, however, there was a marked increase in CAR4^High^ EC, representing almost 20% of total EC, corresponding with the transition from the fetal to neonatal pulmonary circulation. The abundance of the CAR4^High^ EC continued to increase over the subsequent two weeks to reach approximately 30% of total lung EC by P14.

We next explored the effect of chronic hyperoxia^15^, an injury that impairs alveolarization and suppresses capillary EC proliferation, on Cap1 and Cap2 abundance. We quantified Cap2 abundance by FACS and found that hyperoxia increased the relative abundance of CAR4^high^ EC from 25-30% to 35-40% of total EC at both P7 and P14. In contrast, hyperoxia exerted the opposite effect on KIT+ EC, decreasing abundance from 43% to 23% at P7, and from 49% to 27% at P14 (Fig. 1E-F). We confirmed this shift in Cap1 vs. Cap2 abundance by analyzing a separate single cell dataset obtained from neonatal mice exposed to normoxia or hyperoxia for 14 days^16^ (Fig.1G). Consistent with our data, in this data set, hyperoxia increased Cap2 abundance at P7 and P14, and markedly decreased Cap1 abundance at all time points. Taken together, these studies demonstrate that Cap2 abundance markedly increase after birth, and hyperoxia exerts opposing effects on the two capillary subtypes to further increase Cap2 abundance.

### Genetic lineage tracing demonstrates that expanded CAP2 in hyperoxia are derived from Tbx2+ EC existing at birth

Although Car4 expression is higher in Cap2 vs. Cap1, a subset of Cap1 express lower levels of CAR4 (Fig. 1A), making FACS separation challenging based upon CAR4 expression alone. To overcome this challenge and to better identify and study Cap2 EC in the immature lung, we took advantage of their unique expression of *Tbx2* to create a novel mouse model that allowed us to genetically trace *Tbx2*+ EC during development and disease. We utilized transgenic mice (*Tbx2*-iCre) that express both green fluorescent protein (GFP) and tamoxifen-inducible Cre (CreERT2) under control of the Tbx2 promoter (Fig. 2A), such that any cells expressing *Tbx2* will also express GFP. Using FACS to separate CD31^+^ EC from other lineages at P7, we showed that the majority of GFP^+^ EC were KIT^-^ (Cap2 EC), representing approximately 28% of total EC. In contrast, 30% of EC were KIT^+^ EC, representing predominantly Cap1 (Fig. 2B). Of note, we did identify a small population of GFP^+^KIT^+^ double positive EC present at P7.

**Figure 2:**
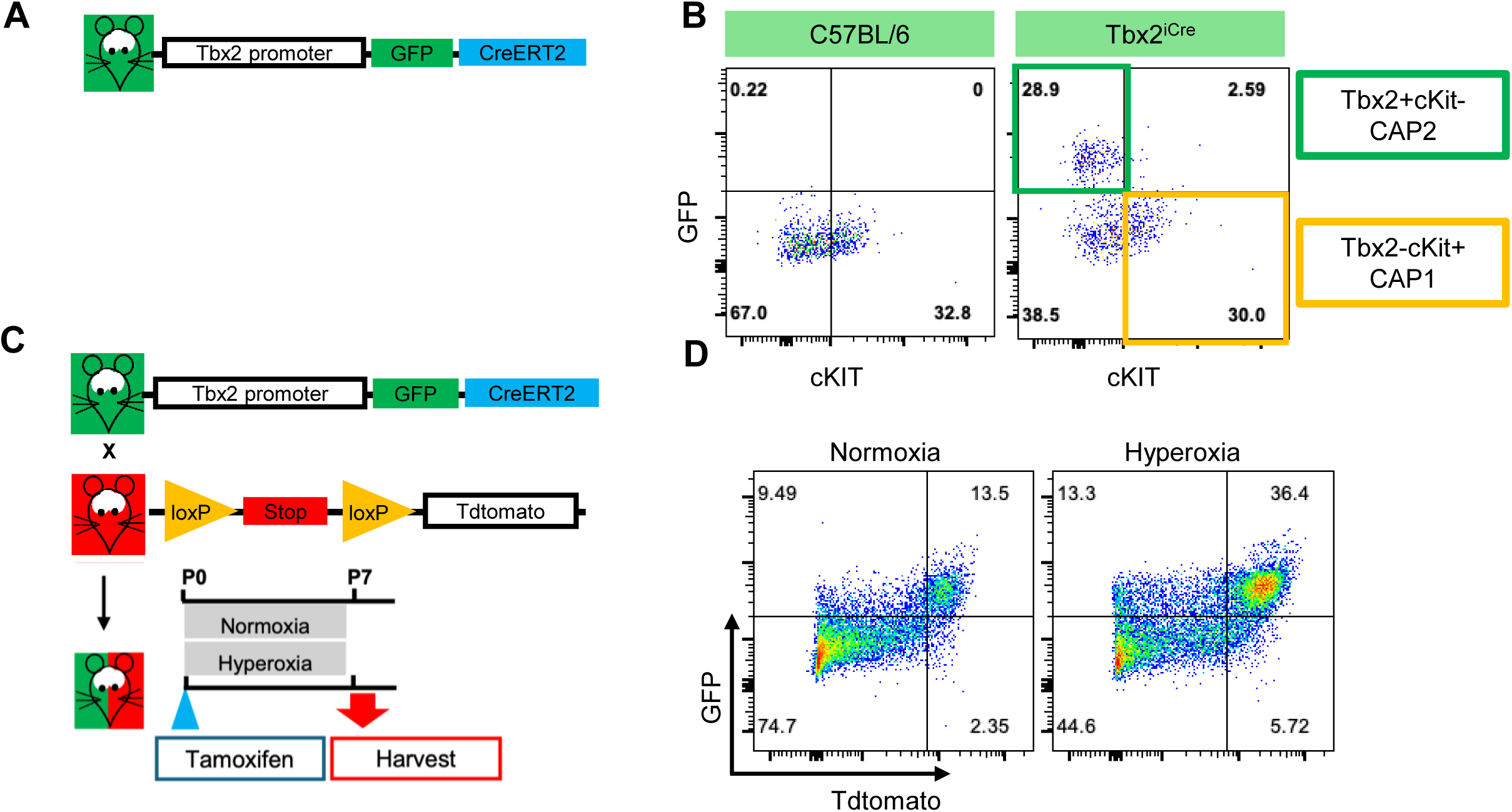
Genetic lineage tracing demonstrates that expanded CAP2 in hyperoxia are derived from Tbx2+ EC existing at birth. **(A)** Schematic of the transgene expressed by the Tbx2-iCre mice. **(B)** Representative FACS plot showing the separation of CD31+ EC by levels of cKIT (x-axis) or GFP (y-axis) expression. **(C)** Schematic of the transgenic constructs used to generate the Tbx2-tdT reporter mice. **(D)** Representative FACS plot showing separation of CD31+ EC by levels of tdT (x-axis) and GFP (y-axis) expression.

The expansion of Cap2 EC after birth and in response to hyperoxia raised the question as to the source of the ‘new’ Cap2 in these settings. In the adult lung, elastase-mediated acute injury does not induce Cap2 proliferation, and recovery appears to result from Cap1 that differentiate into new Cap2. In contrast, proliferating EC that promote vascular regeneration arise from multiple microvascular EC populations^18^. Early alveolarization is marked by rapid growth of the alveolar capillary network, potentially conferring greater cellular plasticity. Thus, we performed genetic lineage tracing of Tbx2^+^ EC, by crossing *Tbx2*-iCre mice with Ai14 reporter mice. In these mice, tamoxifen (TM) will induce tdT in all *Tbx2*-expressing cells at the time of administration, and the tdT lineage label will be retained even if cells undergo subsequent de-differentiation. Importantly, the Tbx2-driven GFP is expressed whenever *Tbx2* is expressed and does not require TM-mediated removal of upstream STOP codon. Thus, this model permits the discrimination of Cap2 that derive from pre-existing Tbx2+ EC (GFP^+^tdT^+^) from those that differentiate later from Cap1 progenitors (GFP^+^tdT^-^) (Fig.2C). In pilot studies, we found that a single dose of 100 ug at P0 labeled approximately 70-80% of Tbx2^+^GFP^+^ EC at P1 (See Figure S1). Subsequently, we administered 100 ug of TM at P0 and exposed mice to normoxia or hyperoxia. At P7, we quantified the percentage of tdT^+^GFP^+^EC by FACS and found that approximately 78% of the GFP+ EC were tdT^+^ (Fig. 2D). Taken together, these preliminary data suggest that the expanded Cap2 in hyperoxia were primarily derived from Tbx2^+^ EC that were present at P0.

### Identification of an intermediate capillary cell state that is present in the developing mouse and human lung

The results of these lineage tracing studies were surprising, as they conflicted not only with the prior reports from the adult lung, but also with a similar study showing that lineage labeling using Kit-Cre^ERT2^ mice also labeled the expanded Cap2 population in hyperoxia^19^. Recalling the small number of Tbx2^+^Kit^+^ double positive EC identified on our FACS analysis (Fig. 2B), we hypothesized that there may be a small progenitor population present during early postnatal development that express both markers. First, we noted within the microvascular EC present at P0, there was a separate Leiden cluster within the Cap1 that expressed lower levels of several Cap2 markers including *Tbx2* and *Car4* (Fig. 3A-B). We also interrogated a single cell data set we recently generated from the late saccular lung that employed endothelial enrichment to ensure the detection of small populations, combined with deep sequencing^20^. These data identified a clear cluster of EC that co-expressed both Cap1 (*Aplnr*, *Plvap*, and *Kit*) and Cap2 (*Tbx2*, *Apln*, and *Car4*) markers and embedded between Cap1 and Cap2 (Fig. 3C-D). We validated the presence of these Cap^INT^ in situ in the P3 lung by performing FISH to detect a combination of Cap1 and Cap2 markers (Fig. 3E). Using this method, we were able to detect Cap^INT^ that expressed all three markers (*Tbx2*, *Kit*, *Pvlap*) at high levels and distinguish them from Cap1 EC that only expressed *Kit* and *Plvap*.

**Figure 3:**
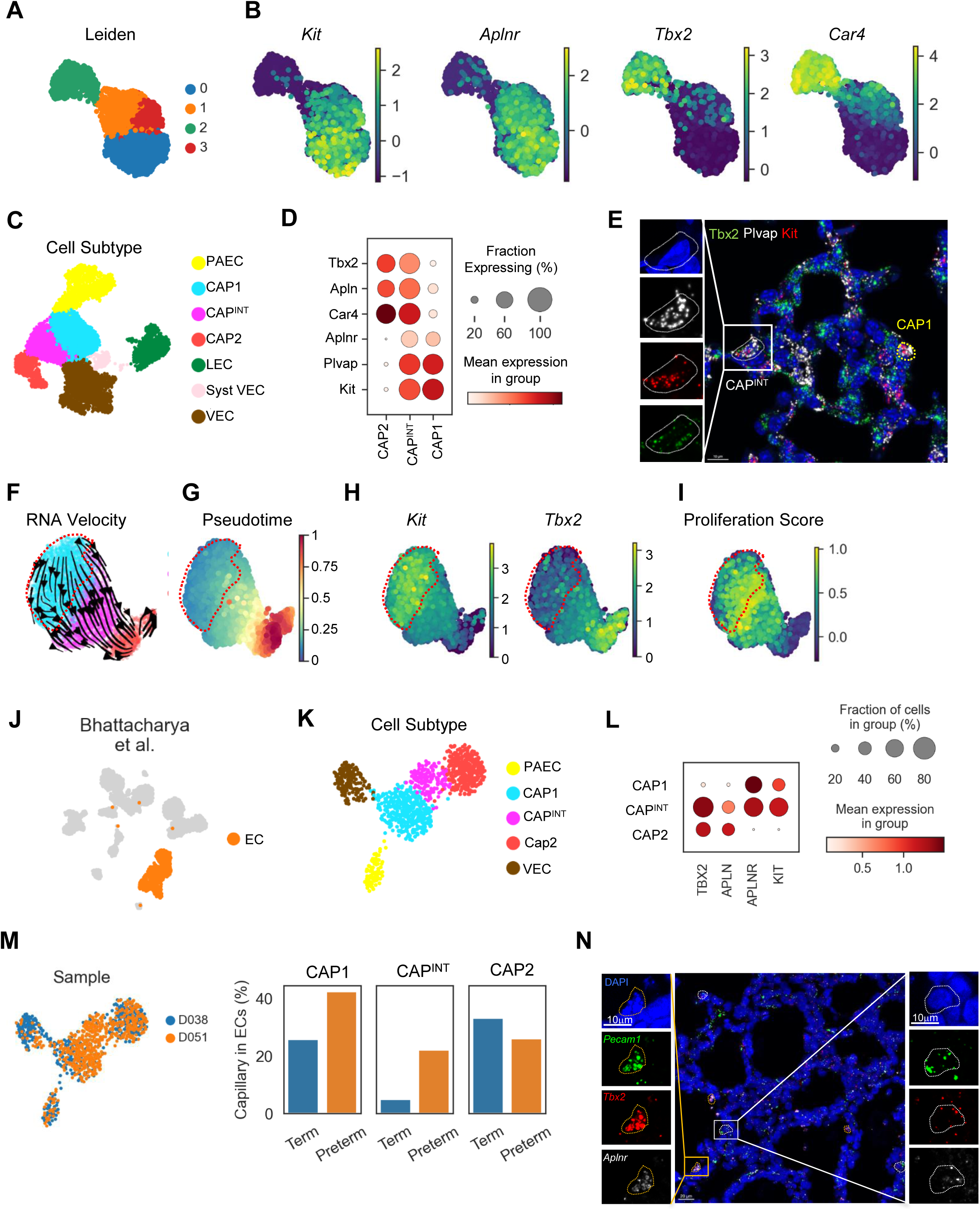
Identification of a novel intermediate capillary cell state that is present in the developing mouse and human Lung. **(A)** UMAP showing re-clustered capillary EC at P0 from the Negretti dataset^17^ by Leiden clusters, and **(B)** feature plots of select Cap 1 and Cap2 marker gene expression showing co-expression within Leiden cluster 1. **(C)** UMAP embedding of EC derived from the lung at P3 from the Sveiven et al. dataset identifying endothelial cell type. PAEC refers to pulmonary arterial endothelial cells, LEC to lymphatic endothelial cells, Syst VEC to systemic vascular endothelial cells, and VEC to venous endothelial cells. **(D)** Dotplot showing expression of Cap1 and Cap2 marker genes in Cap2, Cap^Int^, and Cap1 EC at P3. **(E)** Representative image of multiplex fluorescent in situ hybridization to detect *Tbx2* (green), *Plvap* (white), and *Kit* (red). High magnification of a select Cap^INT^ expressing all three marker genes in contrast to a Cap1 EC outlined with the yellow dotted line. Scale bar=10μm. **(F)** RNA velocity vectors demonstrating a trajectory from Cap1 EC through the Cap^INT^ state to Cap2 EC. **(G)** Embedding of Cap EC along diffusion based pseudotime and **(H)** UMAPs showing the gradual decrease of *Kit* and increase of *Tbx2* along pseudotime. Red dotted circles delineate the Cap1 cluster (**I**) UMAP identifying cells with high proliferation score (yellow) localizing to the late Cap1 and early Cap^INT^ cell state. **(J)** Human lung data re-analyzed from Bhattacharya et al. identifying lung EC, and **(K)** establishing endothelial cell subtyping. **(L)** Dotplot showing expression of Cap1 and Cap2 marker genes in Cap2, Cap^Int^, and Cap1 EC at DOL1. **(M)** Cap EC UMAP embedding visualized by human donor, with D038 a term infant donor, and D051 a donor of 31-week gestation, and quantification of each capillary subtype by donor. **(N)** Representative image of multiplex fluorescent in situ hybridization to detect *Pecam1* (green), *Tbx2* (red), and *Aplnr* (white). High magnification of a select Cap^INT^ expressing all three marker genes (right; white), and (left; orange) a second phenotype of Cap^INT^ exhibiting nuclear condensationand high expression of the marker genes.

Next, we performed trajectory analysis on the capillary EC using RNA velocity (Fig. 3F) to determine whether the Cap^INT^ served as the progenitor population for both Cap1 and Cap2 EC. These data suggest that rather than the Cap^INT^ serving as the progenitor population, a subset of Cap1 exhibited a trajectory through the Cap^INT^ to the Cap2 cell state. Based on that data, we ordered the cells by pseudotime using a single Cap1 EC as the ‘root’ cell (Fig. 3G) which demonstrated that Cap1 canonical markers gradually decrease with advancing pseudotime while Cap2 marker genes increase (Fig. 3H). Utilizing a score calculated from the expression of cell cycle genes we found that highly proliferative capillaries tended to be found at pseudotimes corresponding to the transition from the Cap1 to Cap^INT^ cell state (Fig. 3I).

Finally, we assessed whether this intermediate capillary EC state is present in the human lung. We analyzed a publicly available sc-RNA-seq dataset generated from two human neonatal lung donors: one term infant and one preterm infant delivered at 31 weeks of gestation, both of whom died on the first day of life^21^. In the original analysis, EC subtypes were not defined (Fig. 3J). To refine the annotation, we re-embedded the EC alone, enabling us to resolve known vascular EC subtypes, including arterial, venous, and both Cap1 and Cap2 EC (Fig. 3K). Using the Leiden clustering algorithm, we identified a distinct cluster situated between the Cap1 and Cap2 populations that co-expressed markers of both subtypes, leading us to annotate it as Cap^INT^ (Fig. 3K-L). Notably, analysis of donor-specific representation revealed that the majority of Cap^INT^ EC originated from the preterm donor, representing more than 20% of the total EC from that donor (Fig. 3M). We validate the presence of these cells in situ using lung tissue from a separate preterm donor who died on DOL#1 (Fig. 3N). We identified several *Pecam1*+ EC expressing a combination of Cap1 and Cap2 markers (*Tbx2* and *Aplnr)*. Of note, there was a separate population of *Tbx2*+*Aplnr*+ EC (orange dotted circle) that demonstrated nuclear condensation with overlapping high expression of the two markers, consistent with actively dividing cells. Taken together, our data identify for the first time, an intermediate alveolar capillary cell state present soon after birth in both the developing human and mouse lung.

### Hyperoxia increases the abundance of Cap^INT^ endothelial cells

We next determined whether hyperoxia altered the balance between Cap1, Cap^INT^, and Cap2 EC during postnatal development. Using the Sveiven et al. dataset from the late saccular lung (P3)^20^, we found that hyperoxia induced a slight increase in Cap^INT^ abundance by this early timepoint (Fig. 4A), which we confirmed by quantifying Cap^INT^ in situ using FISH (Fig. 4B). To validate these findings, we also re-analyzed the hyperoxia data set generated by Hurskainen et al. which included later hyperoxia-exposed timepoints. In this analysis, we re-clustered the capillary EC, and again identified clusters of Cap^INT^ EC embedding between Cap1 and Cap2 EC (Fig. 4C). Quantifying each capillary cell subtype in each condition at each timepoint (Fig. 4D) showed that Cap2 abundance continued to increase across the 14-day hyperoxia exposure. Concurrently, Cap1 abundance decreased with a marked reduction in Cap1 in P14 pups exposed to chronic hyperoxia. Consistent with our data sets, Cap^INT^ represented approximately 10% of total EC at P3 but were rare in the normoxic lung at later developmental timepoints. In contrast, hyperoxia increased Cap^INT^ abundance such that these EC represented approximately 20% of total EC at P7 and P14.

**Figure 4:**
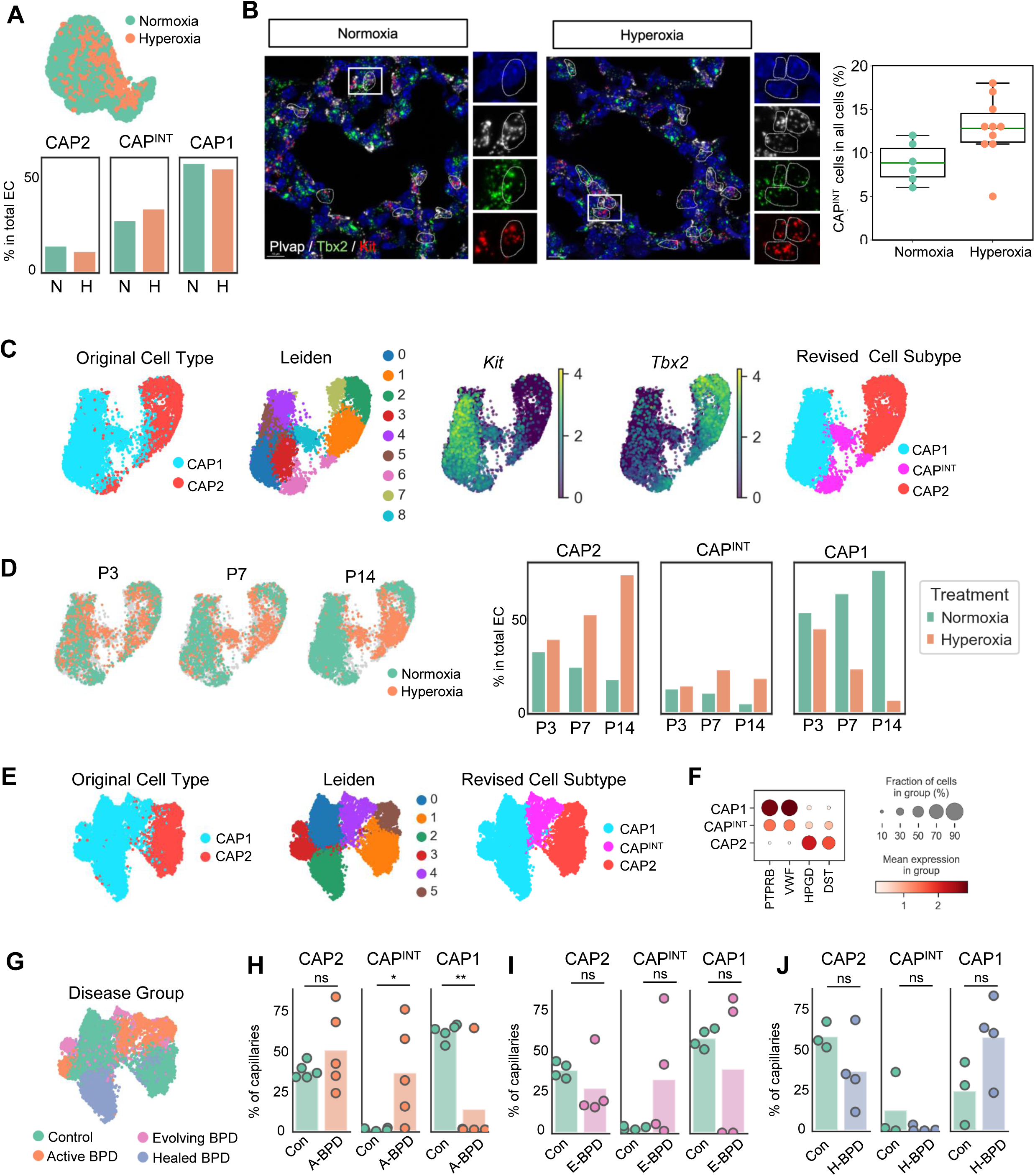
Hyperoxia Increases the Abundance of Cap^INT^ Endothelial Cells. **(A)** UMAP embedding of capillary EC from the murine lung at P3 from the Sveiven et al. dataset visualizing cells from each treatment with quantification of abundance as a percentage of total EC. **(B)** Representative images of multiplex fluorescent in situ hybridization to detect *Tbx2* (green), *Plvap* (white), and *Kit* (red) in mouse lungs at P3 exposed to either normoxia or hyperoxia, with quantification of Cap^INT^ EC in each condition as compared to total lung cells. Each dot represents a selected area used for quantification; Data are presented as mean ± SEM from n = 2 mice per group **(C)** UMAP embedding of capillary EC from Hurskainen et al. visualizing original cell typing, Leiden clustering, *Kit* and *Tbx2* expression and revised cell subtyping. **(D)** UMAP visualization of data from Hurskainen et al. identifying EC from each experimental group at each timepoint with quantification of each capillary EC subtype as a percentage of total lung EC. **(E)** Human data obtained from LungMap (LMEX0000004400) with UMAP embeddings showing original cell type annotation, new Leiden clusters, and revised cell type annotation. **(F)** Dot plot showing the expression of Cap1 and Cap2 marker genes in human lung. **(G)** UMAP embedding visualizing EC derived from each clinical group. **(H)** Quantification of each cap EC subtype in each clinical treatment group as a percentage of capillaries, (n=3-5), each point represents a sample, and the bar represents the mean of each group, student’s t-test was run between experimental groups versus age-matched controls, with *=*p*<0.05, **=*p*<0.01

We next explored whether alterations in these capillary fate transitions occur in infants who develop bronchopulmonary dysplasia (BPD), a chronic lung disease marked by impaired alveolarization and distal lung angiogenesis. We obtained single cell data from LungMap (LMEX0000004400) from cadaveric samples corresponding to four patient groups. Initial cell typing was then revised based on Leiden clustering and the co-expression of Cap1 and Cap2 marker genes detected in this dataset (Fig. 4E-F). Patients with active BPD had a diagnosis of severe BPD and died from respiratory failure, patients with evolving BPD were still receiving respiratory support but died from other causes, patients with ‘healed’ BPD were no longer receiving respiratory support and age-matched controls for each group. Visualizing cell type distribution by disease group showed that Cap^INT^ were only found in patients with active and evolving BPD (Fig. 4G). In patients dying from active BPD, there was no difference in Cap2 EC abundance, however Cap^INT^ were markedly increased and Cap1 decreased (Fig. 4H). In patients with evolving BPD, Cap^INT^ abundance remained elevated and Cap1 EC remained decreased but did not reach statistical significance given the small sample size. Finally, in healed BPD, Cap^INT^ were rare in both patients and controls, and there were no significant differences between Cap2 and Cap1 abundance.

Taken together, our data identify an intermediate cell state between Cap1 and Cap2 EC in both the developing human and murine lung. Cap^INT^ cells are abundant in the lung after birth but decrease in abundance as alveolarization progresses. However, hyperoxia increases Cap^INT^ abundance in the developing lung, and proliferating EC are enriched as cells transition from the late Cap1 to the early Cap^INT^ cell state. Alterations in capillary cell type abundance is apparent in patients with bronchopulmonary dysplasia with increases in Cap^INT^ EC present in patients with active BPD.

### Postnatal loss of VEGFR2 in Tbx2+ EC increases Cap^INT^ abundance and blocks Cap2 terminal differentiation

Prior studies have shown mice lacking epithelial-derived *Vegfa* lack Cap2 and exhibit impaired alveolarization. However, whether VEGF signaling is required postnatally to maintain Cap2 identify, and the VEGFR2 downstream mechanisms that regulate lung capillary EC subtype fate decisions are not know. To address these gaps in knowledge, we created a novel mouse model to allow selective, abrogation of VEGFR2 in Tbx2+ EC during key timepoints in development by breeding *Tbx2*-GFP-iCre mice to Kdr-floxed mice (the gene encoding VEGFR2). In pilot studies, we administered TM from P0-P2 and confirmed by FACS that this strategy decreased VEGFR2 protein expression, decreasing the number of VEGFR2^HIGH^ EC by P7 to 26% (compared to 92% of EC in WT mice) (Fig. 5A). We confirmed in VEGFR2^LOW^ EC isolated from the KO mice that this strategy effectively deleted exon3 of *Kdr* but did not induce compensatory alterations in *Flt-1*, the gene encoding VEGFR1 (Fig. 5B). Next, we assessed whether inducible loss of VEGFR2 altered Cap2 abundance at baseline or the expansion of Cap2 EC in response to hyperoxia. Tbx2-iCre-Kdr^WT/WT^ (VEGFR2^WT^) and Tbx2-iCre-Kdr^f/f^ (VEGFR2^ΔCap2^) mice were administered daily TM from P0-P2 and exposed to normoxia or hyperoxia. In normoxic mice, the abundance of GFP positive EC at P7 was not different between genotypes (25.3% vs. 24.2%), although the level of GFP expression appeared decreased in the KO mice (Fig. 5C). Hyperoxia increased GFP+ EC from 25% to 42% in VEGFR2^WT^ mice, however, this increase was blocked in VEGFR2^ΔCap2^ mice.

**Figure 5:**
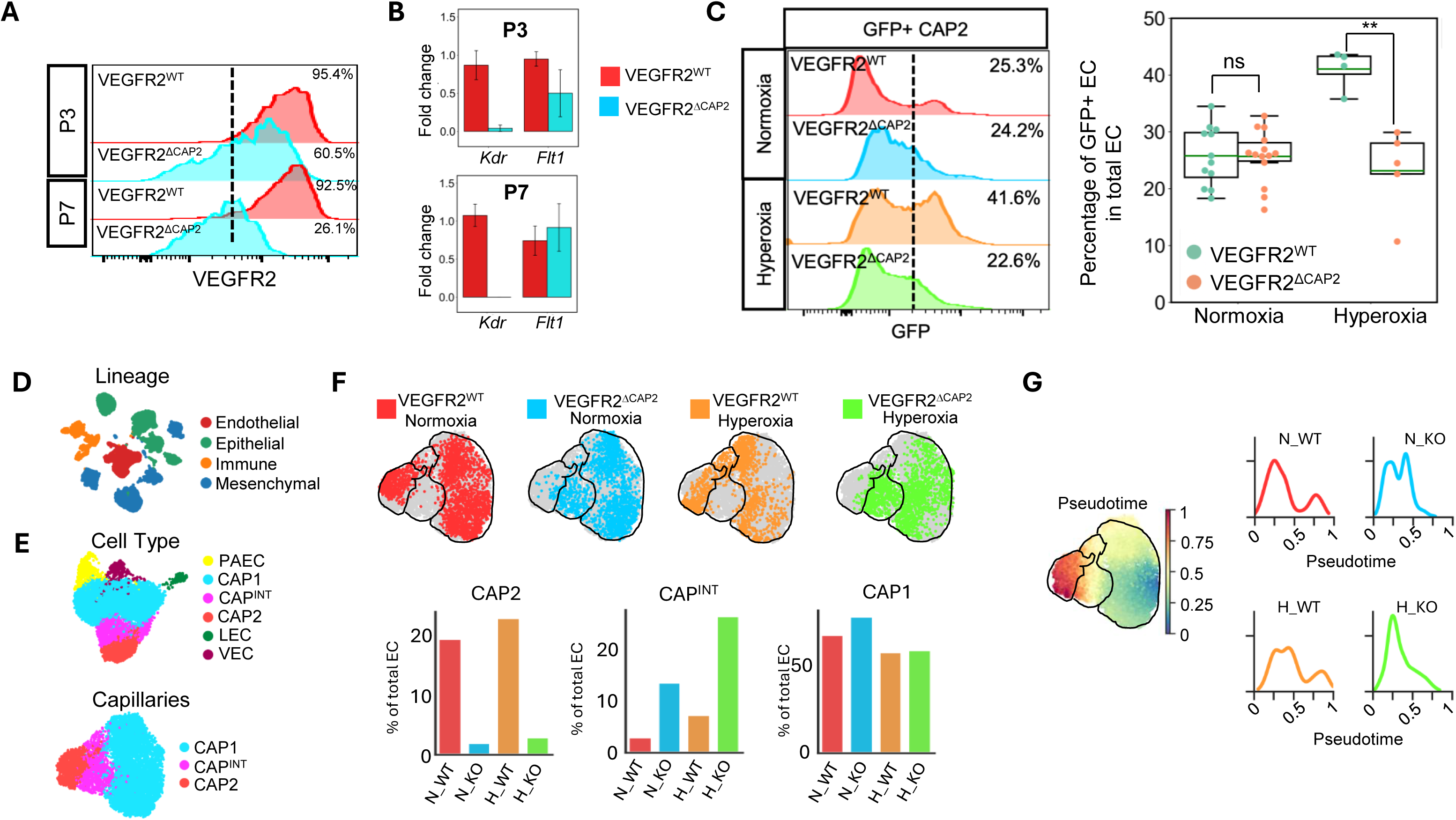
Postnatal Loss of VEGFR2 in Tbx2+ EC Increases Cap^INT^ Abundance and Blocks Cap2 Terminal Differentiation. **(A)** Representative FACS plot to assess VEGFR2 protein expression in VEGFR2^WT^ and VEGFR2^ΔCAP2^ mice maintained in normoxia or hyperoxia at P3 and P7. **(B)** qPCR to specifically detect exon 3 of the Kdr gene and Flt-1 normalized to β-actin. Data are presented as mean fold change ± SEM relative to control, with n=2-3 mice per group **(C)** Representative FACS plot and quantification to assess GFP protein expression in VEGFR2^WT^ and VEGFR2^ΔCAP2^ mice maintained in normoxia or hyperoxia at P7 with n=4-15 per group. Data are presented as mean ± SEM **p=0.003 via unpaired two-tailed Student’s *t*-test. **(D)** UMAP plot of transcriptomic data from 31,432 single nuclei demonstrating representation from all major lineages in the lung. **(E)** Separate embeddings of 7,353 lung EC (top), including 6,733 capillary EC (bottom). **(F)** UMAP visualization of capillary cells of each subtype in each condition and genotype, with quantification as a percentage of total EC. **(G)** Embedding of cells visualized by pseduotime with the plots showing the distribution of capillary ECs across pseudotime value in each experimental condition and genotype.

To better define how loss of VEGFR2 alters capillary subtype identity, we performed sn-RNA-seq on 31,432 lung cells from VEGFR2^WT^ and VEGFR2^ΔCap2^ mice at P7 from each treatment group (Fig. 5D). We then separately analyzed the 7,353 vascular EC to obtain better resolution to detect differences between the two genotypes (Fig. 5E). All three capillary EC subtypes were represented, including a large cluster of Cap^INT^ EC. Visualization by genotype and treatment (Fig. 5F) identified clusters that were derived predominantly from either VEGFR2^WT^ or VEGFR2^ΔCap2^ mice in normoxia or hyperoxia. We quantified capillary subtype abundance and found in normoxic VEGFR2^WT^ mice (Fig. 5F, red bars), that the majority of EC were Cap1, with Cap^INT^ representing only 3.7% of total EC, consistent with our prior findings. Hyperoxia increased Cap^INT^ abundance 3-fold in the VEGFR2^WT^ mice, and slightly increased Cap2 EC (orange bars). In contrast, in the VEGFR2^ΔCap2^ mice (blue and green bars), Cap2 EC were almost completely absent, and Cap^INT^ abundance increased to 16% of total EC in normoxic mice and 30% of total EC in hyperoxic mice. To better delineate these fate transitions that appear to exist along a continuum, we ordered the cells along pseudotime, using a single Cap1 EC as the root cell. In WT mice maintained in normoxia, EC were predominantly found at either low or high pseudotime, with very few EC found at intermediate pseudotime (consistent with the Cap^INT^ cell state). Hyperoxia pushed this curve rightward in the WT animals, increasing the number of EC located between pseudotime values of 0.25 and 0.5. This effect was much more dramatic in the KO mice, where in normoxia, there was a stark absence of EC at the latest pseudotime (corresponding to the most differentiated Cap2 EC) and markedly increasing EC with intermediate pseudotime values, with a further right ward shift of this curve in hyperoxia exposed KO mice. Taken together, these studies demonstrate that VEGFR2 is not required for entry into the Cap^INT^ cell state but is indispensable for terminal differentiation of Cap2 after birth.

### Postnatal abrogation of Cap2 differentiation disrupts alveolar formation and induces α-SMA expression in the distal Lung

To understand how these alterations in capillary subtype abundance and phenotype affect the structure of the alveolar niche, we performed histologic analysis of alveolar structure in the VEGFR2^WT^ or VEGFR2^ΔCap2^ mice at baseline and in response to chronic hyperoxia. We first performed immunofluorescent staining of thick vibratome sections of mice at P7 to detect CD31, the AT1 specific marker, AQP5, and α-smooth muscle actin (Fig. 6A). Loss of VEGFR2 in Cap2 resulted in areas with obvious airspace dilation (asterisk). Further, even in areas where alveolar size was similar to controls (dotted line encircled area, and high magnification inset on left), there were alterations in capillary network formation. In contrast to almost complete coverage of the airspace with tiny ‘pores’ in the WT mice, the capillary network of the VEGFR2^ΔCap2^ mice appeared ‘moth-eaten’. Moreover, there was a strong increase in α-SMA along the alveolar rings in VEGFR2^ΔCap2^ mice.

**Figure 6:**
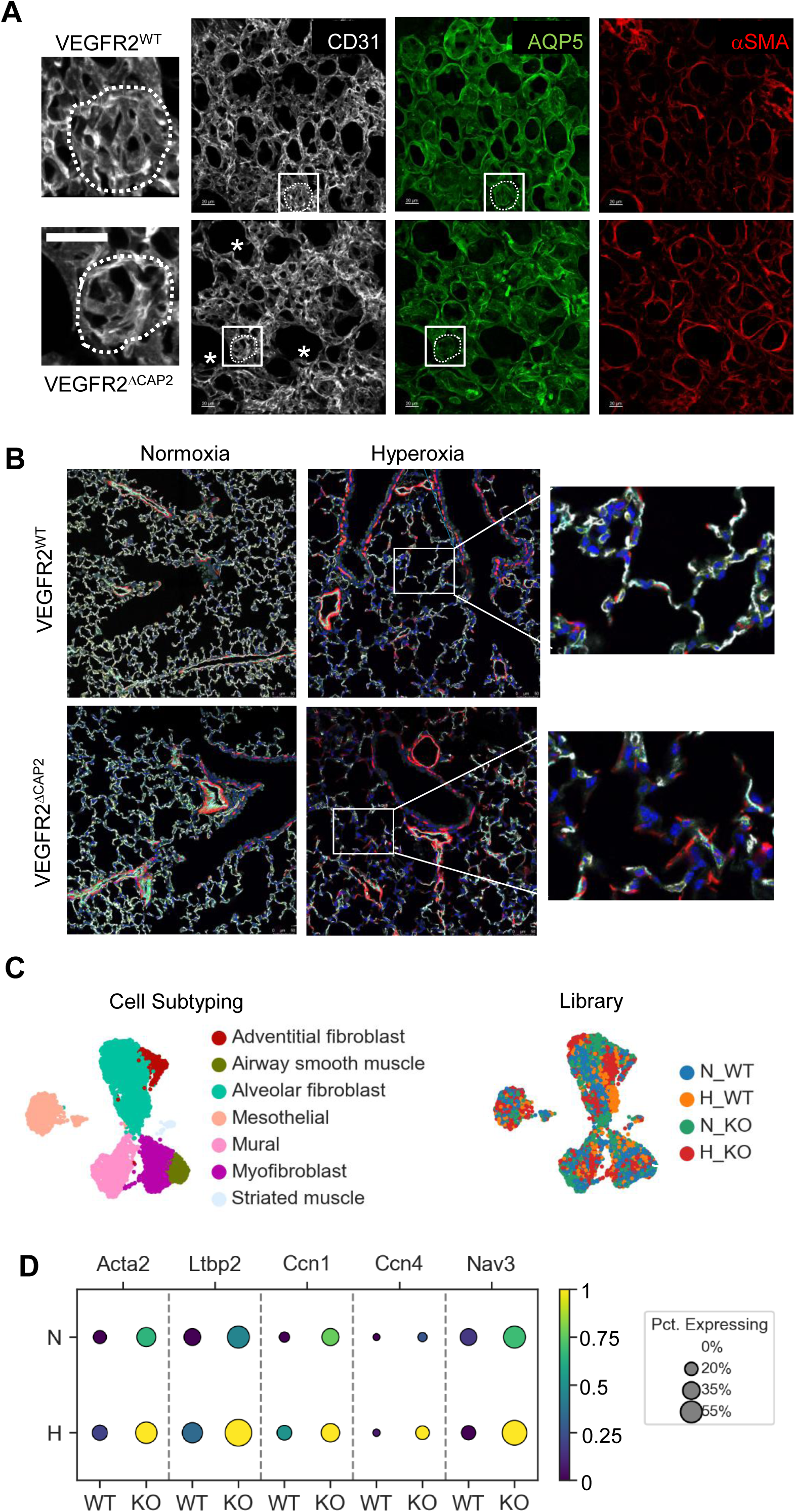
Postnatal Abrogation of Cap2 Differentiation Disrupts Alveolar Formation and Induces α-SMA Expression in the Distal Lung. **(A)** Representative immunofluorescent images of vibratome sections of lung from normoxia-exposed VEGFR2^WT^ and VEGFR2^ΔCAP2^ mice stained to detect CD31 (white), AQP5 (green) and α-SMA (red) at P7. Asterisk denotes enlarged airspaces. High magnification inset identifies differences in the microvascular structure surrounding alveoli of similar size. Calibration mark=20 μm. **(B)** Representative immunofluorescent images of lung sections from normoxia- and hyperoxia-exposed VEGFR2^WT^ and VEGFR2^ΔCAP2^ mice stained to detect CD31 (white) and α-SMA (red) at P14. High magnification insets showing alterations in EC coverage and α-SMA+ cells in the distal lung. **(C)** UMAP of single nuclei data from mesenchymal cells demonstrating representation of all mesenchymal cell subtypes (left) and visualized by experimental group (right). **(D)** Dotplot showing altered expression of numerous genes involved in fibroblast proliferation, migration, and activation.

We also performed similar studies to assess alterations in lung structure between the two genotypes at a later timepoint (P14) in groups maintained in normoxia or exposed to hyperoxia (Fig. 6B). Compared to VEGFR2^WT^ mice, distal airspace dilation was still apparent in VEGFR2^ΔCap2^ mice under control conditions. Hyperoxia induced marked airspace dilation in both genotypes. However, the dilated distal airspaces in VEGFR2^WT^ mice exhibited continuous coverage by CD31+ EC, and limited expression of α-SMA (high magnification inset, top). In contrast, the distal airspaces of the hyperoxia-exposed VEGFR2^ΔCap2^ mice exhibited discontinuous endothelial coverage, with many areas completely devoid of endothelial cells. Of note, in many of the surfaces lacking CD31+ EC, there was an increase in α-SMA+ cells. We confirmed this increase in α-SMA expression in the lung by WB which confirmed an increase in α-SMA expression in the VEGFR2^ΔCap2^ mice that was further exaggerated by hyperoxia exposure (See Figure S2).

These alterations in lung structure at this early timepoint suggested that loss of terminally differentiated Cap2, or persistence of aberrant Cap^INT^ may promote maladaptive angiocrine signaling to induce non-cell-autonomous alterations in the phenotype of other lung cells to further impair development and increase fibrosis. This latter hypothesis is reminiscent of the paradigm established in the alveolar epithelium, where AT2 enter a transition state before differentiating into AT1 cells in development and physiologic repair, but in chronic injury persistence of aberrant AT1-2 intermediates promote lung fibrosis^22,23^. Thus, we next explored transcriptomic alterations in the mesenchymal cells within the alveolar niche that might be responsible for the increased α-SMA expression apparent in the VEGFR2^ΔCap2^ mice. We profiled 10,989 mesenchymal cells, with representation of all major subtypes and cells in each subtype derived from each library (Fig. 6C).

Of the cells present in the alveolar niche, only alveolar fibroblasts (AF) demonstrated an increased expression of *Acta2*, encoding α-SMA. Moreover, AF from the VEGFR2^ΔCap2^ mice exposed to hyperoxia exhibited an increase in numerous genes associated with activation and transition to a contractile, myofibroblast like phenotype. These genes include a marked up-regulation in *Ltbp2*, a TGFβ binding protein increased in activated fibroblasts in pulmonary fibrosis^24^, *Cnn1* and *Cnn4*, genes encoding matricellular proteins that promotes fibroblast activation that are increased in AF isolated from patients with progressive lung fibrosis^25^, and *Nav3*, a sodium channel that promotes fibroblast migration and contractility, increased in kidney fibrosis^26^ (Fig. 6D).

In summary, our data demonstrate that postnatal loss of VEGFR2 induces airspace dilation and disrupts the alveolar capillary network at baseline and induces loss of endothelial coverage of the airspaces and α-SMA + cells expression in the distal lung in response to hyperoxia. These structural changes were associated with non-cell autonomous effects on alveolar fibroblasts, which display a transcriptomic signature consistent with activation. This data suggests that terminally differentiated Cap2 are essential for normal alveolarization and maintenance of a homeostatic alveolar capillary niche.

### Hyperoxia increases ERG expression and chromatin accessibility which is lost by postnatal loss of VEGFR2

Finally, we aimed to identify mechanisms downstream from VEGFR2 that are required for Cap2 terminal differentiation. Given that hyperoxia pushes capillary progenitors toward the Cap2 fate, we hypothesized that interrogation of differential chromatin accessibility in Cap1 versus Cap2 in hyperoxia versus normoxia may identify transcriptional regulators driving Cap2 differentiation. FACS quantification showed that hyperoxia-mediated increases in Cap2 abundance began after P3, with marked expansion observed by P7 (Fig. 7A). Thus, to explore transcriptional regulators that promote Cap2 differentiation, we selected P3 as a key timepoint for analysis. We used Tbx2-driven GFP expression to FACS separate GFP^HIGH^ (Cap2 EC) from GFP^LOW^ EC (non-Cap2 EC) from P3 mice exposed to normoxia or hyperoxia and subjected the four groups to bulk ATAC-seq (Fig. 7B). Principal component analysis (PCA) revealed differences in chromatin accessibility, with samples clustering based on cell type (PC-1) and treatment (PC-2) (Fig. 7C).

**Figure 7:**
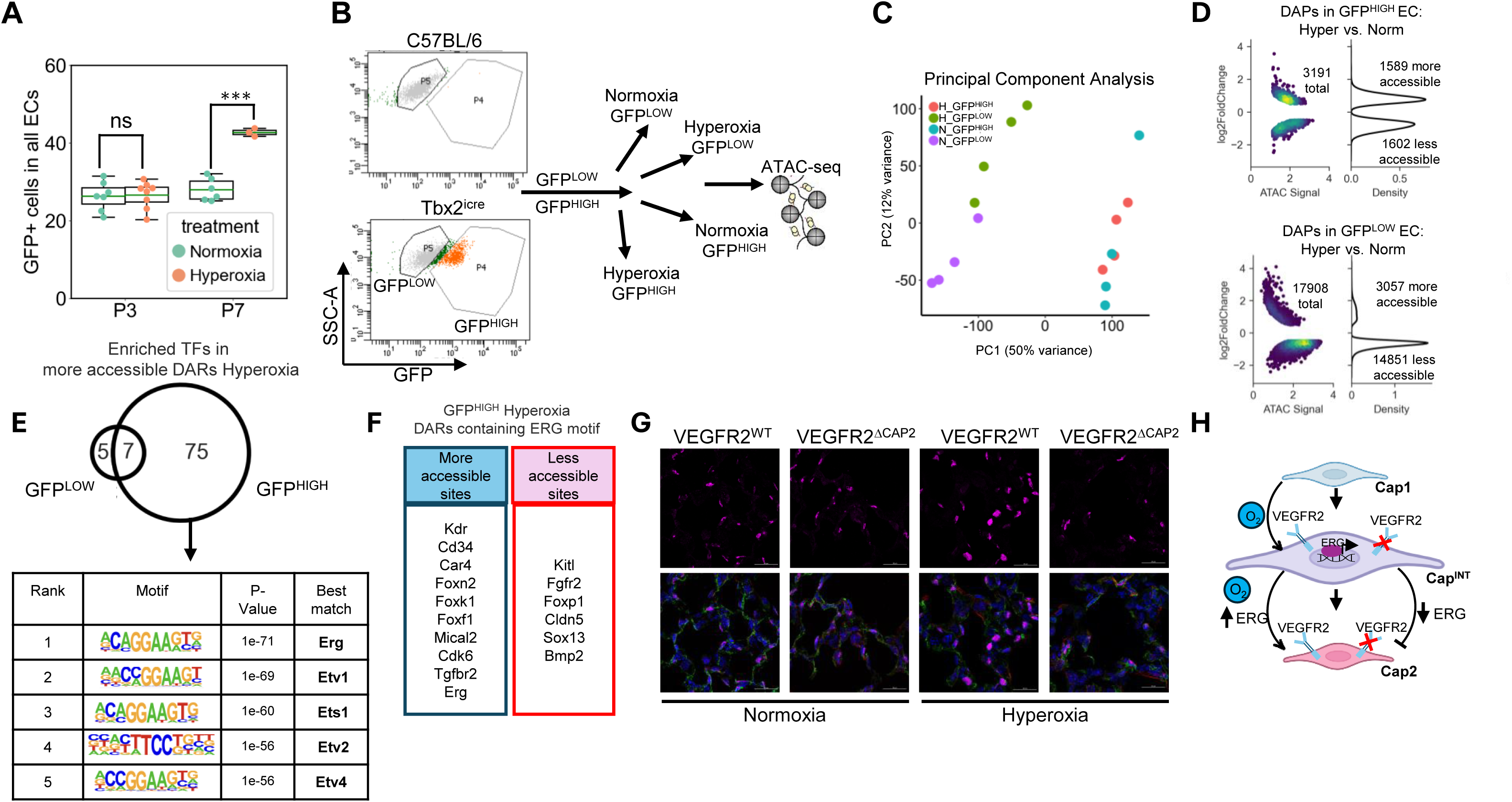
Hyperoxia Increases ERG Expression and Chromatin Accessibility which is Lost by Postnatal Loss of VEGFR2. **(A)** FACS data quantifying the percentage of GFP+ EC in total EC in mice exposed to normoxia or hyperoxia at P3 and P7. Data are mean ± SEM, with n=3-8 mice per group and *** *P* =0.00008. **(B)** Representative FACS plot showing the gating strategy to isolate GFP low and high EC from normoxia and hyperoxia exposed mice at P3 for ATAC-seq. **(C)** PCA plot of the four experimental groups with samples separating by cell type and treatment. **(D)** Kernel density estimates plots showing the number of differentially accessible peaks (DAP) in hyperoxic versus normoxic GFP^HIGH^ (top) and GFP^LOW^ (bottom) EC. The density of reads corresponding to a particular peak is visualized as low (purple) to high (yellow) on the left plot and the distribution right hand plot. **(E)** Venn diagram showing overlap of enriched transcription factors with sites in DAPs more accessible in hyperoxia in GFP^HIGH^ or GFP^LOW^ ECs **(F)** Table including the top predicted enriched TF binding domains in the DAPs in the hyperoxic versus normoxic GFP^HIGH^ EC. **(G)** Table including select genes with ERG binding motifs annotated to the TSS of DAPs in hyperoxic versus normoxic GFP^HIGH^ EC. **(H)** Representative IF to detect ERG (purple) in combination with DAPI (blue) and CD31 (green) and α-SMA (red) in P7 VEGFR2^WT^ and VEGFR2^ΔCAP2^ mice exposed to normoxia or hyperoxia. Cal bar=20μm. **(I)** Proposed model.

We next aimed to identify transcriptional factors (TFs) promoting Cap2 expansion. We performed de novo motif analysis within differentially accessible peaks (DAPs) in hyperoxic GFP^HIGH^ versus normoxia, focusing on putative TFs that were not similarly enriched in hyperoxic versus normoxic GFP^LOW^ EC. We compared the number of differentially accessible peaks (DAPs) in the GFP^HIGH^ EC in hyperoxia vs. normoxia, and found over 3000s DAPs, that were evenly split into regions that were either more or less accessible (Fig. 7D). In contrast, in the GFP^LOW^ EC, although there were more DAPs, most of these peaks were less accessible in hyperoxia.

To identify transcriptional factors (TFs) driving Cap2 expansion, we performed de novo motif analysis within differentially accessible peaks (DAPs) in GFP^HIGH^ EC in hyperoxia versus normoxia, and again in GFP^LOW^ EC. Of note, the number of unique enriched TF binding sites in GFP^HIGH^ EC was higher than the number of unique TF binding sites in GFP^LOW^ in DAPs more open in hyperoxia (Fig. 7E). We found that binding motifs for ERG emerged as the most enriched in DAPs in hyperoxic versus normoxic GFP^HIGH^ EC (Fig. 7E), with many other ETS family TFs also found to be enriched. This contrasted with the hyperoxic versus normoxic GFP^LOW^ EC where p53 was the most enriched putative TF in the DAPs (See Figure S3), and enrichment of ERG binding motifs was not found. Next, we annotated the nearest transcriptional start site (TSS) to each DAP. This analysis identified increased accessibility in numerous chromatin regions containing ERG binding motifs in the hyperoxic GFP^HIGH^, including genes associated with Cap2 identity (e.g. *Kdr, Cd34*, *Car4*, *Foxf1* and *Foxk1*). Conversely, genes associated with Cap1 identity (e.g. *Kitl, Cldn5* and *Sox13)* showed less chromatin accessibility within ERG binding regions in hyperoxic Cap2 (Fig. 7F).

Next, we compared the effect of hyperoxia on ERG expression and whether this was altered by postnatal VEGFR2 deletion. We immunostained lung sections to detect ERG from the VEGFR2^WT^ and VEGFR2^ΔCap2^ mice. At baseline, ERG immunoreactivity appeared similar between the two groups. However, in the WT mice, hyperoxia markedly increased the intensity of ERG nuclear staining, but this effect was completely blocked in the VEGFR2^ΔCap2^ mice (Fig. 7G). These data suggest that hyperoxia enhances ERG expression and ERG chromatin accessibility in the Cap2 EC, and that loss of VEGFR2 blocks the hyperoxia-mediated increase in ERG. Given that loss of VEGFR2 in Tbx2+ EC also blocks Cap2 expansion in VEGFR2^ΔCap2^ mice exposed to hyperoxia, highlights ERG as a potential VEGFR2-mediated downstream activator of Cap2 fate (Fig. 7H).

## Discussion

The alveolar capillary network is maintained by two distinct capillary populations, one with proliferative potential to facilitate capillary growth and repair (Cap1), and the other serving a specialized role in gas exchange (Cap2). However, the molecular mechanisms directing capillary speciation, developmental plasticity, and fate transitions during alveolar formation and repair are not well understood. In this study, we show that the Cap2 cells rapidly expand after birth and are further expanded by hyperoxia. We show that expansion of the Cap2 population requires two distinct steps. First, Cap1 progenitors enter an intermediate cell state, with Cap^INT^ EC expressing markers of both subtype and heightened proliferation. These Cap^INT^ EC are present during development in the murine and human lung but decrease in abundance as alveolarization progresses. Hyperoxia increases Cap^INT^ abundance in developing mice, and increased Cap^INT^ are observed in infants with active BPD, a chronic lung disease marked by disrupted alveolarization and angiogenesis. Although VEGRF2 is dispensable for entry into the Cap^INT^ cell state, it is required for terminal differentiation of Cap^INT^ into Cap2, and loss of VEGFR2 in Tbx2^+^ EC results in both an absence of fully differentiated Cap2 EC and a marked increase in Cap^INT^. These changes in cell abundance were associated with structural alterations in the developing lung including airspace dilation and an increase in molecular changes consistent with fibroblast activation. Finally, we identified ERG as a putative mediator downstream of VEGFR2 that appears to play a key role in promoting Cap2 differentiation.

Prior work has shown that loss of epithelial-derived VEGFA during embryonic development blocks Cap2 speciation. Yet, VEGFA treatment alone is not sufficient for Cap2 speciation^14^. Moreover, although mice lacking AT1-derived VEGFA exhibit impaired alveolar formation, it was not clear whether these structural abnormalities resulted from the absence of Cap2 speciation, the overall reduction of VEGFA, or a combination of both defects. VEGFR2 is required for adaptive angiocrine signaling in the adult lung, yet whether the high expression of VEGFR2 in Cap2 heralds a central role for these EC in establishing a homeostatic alveolar capillary niche has not been explored previously. One key finding from our novel mouse model was that postnatal abrogation of VEGFR2 signaling in Cap^INT^ EC and subsequent prevention of Cap2 terminal differentiation was sufficient to impair alveolarization and induce molecular changes consistent with activation of alveolar fibroblasts. These results suggest that abrogating physiologic Cap1-Cap^INT^-Cap2 fate transitions causes non-cell autonomous effects via maladaptive angiocrine signaling. Future research to identify these non-cell-autonomous mechanisms are warranted to provide insight into how dysregulated ECs may promote fibrosis in lung disease models. Given that decreased VEGF signaling has been implicated in numerous lung diseases, including bronchopulmonary dysplasia, ARDS, and COPD^1,27,28^, delineating how this key pathway regulates Cap2 differentiation and function has broad implications for pediatric and adult lung diseases marked by impaired growth, repair and fibrosis.

Our data suggest that ERG is a key VEGFR2 downstream mechanism that promotes Cap2 terminal differentiation in response to hyperoxia. In WT mice, hyperoxia increased Cap2 abundance in association with increased nuclear immunoreactivity for ERG and increased chromatin accessibility of regions containing ERG binding sites. A number of these binding sites were annotated to genes that are canonical Cap2 markers. Loss of VEGFR2 in Tbx2^+^ EC blocked the increase in nuclear ERG protein, further implicating ERG as downstream of VEGFR2 in this model. ERG is a member of the E-26 transformation specific (ETS) TF family, a key regulator of transcriptional programs which promotes EC homeostasis, vascular stability, and angiogenesis^29^. In human microvascular EC, VEGF activates MEK/ERK, leading to ERG phosphorylation. The direct targets of ERG-mediated transcription in the Cap^INT^ remain to be elucidated, as well as knowledge regarding whether ERG promotes Cap2 differentiation alone or does so in concert with addition transcription factors.

The identification of an intermediate cell state in the capillary EC parallels a similar paradigm in alveolar epithelial cells, where alveolar type 2 cells (AT2) transition to an intermediate state prior to differentiation to AT1 cells. During development, both AT1 and AT2 cells appear to arise from bipotent progenitors that co-express marker genes of both cell types^30^. However, in lung injury, AT1 cell death induces AT2 proliferation and transition to damage associated transitional progenitors (DATPs) prior to differentiation into AT1 cells. These DATPs highly express integrin β6, and several pro-fibrotic mediators including transforming growth factor beta that serve to activate fibroblasts to promote lung fibrosis^31^. In a bleomycin model of lung injury, the DATP cell state appears to be driven by inflammatory signals (e.g. Il1β) produced by lung interstitial macrophages, with *Il1r* expressing AT2 cells contributing predominantly to the DAPT cell pool via pathways which require HIF1α signaling. Additional studies in our model will be important to understand if hyperoxia activates resident lung immune cells that serve as a key regulator of the Cap^INT^ state in developmental lung injury.

To our knowledge, this is the first report showing that Cap1 progenitors enter an intermediate capillary cell state prior to differentiation into Cap2 during normal development in both the mouse and human. Our data suggest that Cap^INT^ EC are enriched in proliferating EC, suggesting that proliferation may be a prerequisite for Cap1 to Cap2 differentiation. In our studies, enrichment of EC from the late saccular murine lung allowed profiling of sufficient EC to appreciate the Cap1-Cap^INT^-Cap2 continuum that was key to identification of this cell state in development. Current single cell algorithms strive to ‘cluster’ cells into discrete entities, often resulting in arbitrary binning of cells of intermediate states into one cluster or another, explaining why these Cap^INT^ were missed in prior reports. However, once we recognized this continuum, subsequent identification of Cap^INT^ in less robust mouse and human datasets was possible.

Recent studies have reported various forms of capillary intermediate cells in several diseases. For example, in adult mice exposed to influenza-induced lung injury, injury-induced (iCap) emerge that also appear to express an intermediate signature. Of note, lineage tracing studies using either *Kit* or *Car4* suggested that these iCap could derive from either Cap1 or Cap2 EC. However, whether a small number of Cap^INT^ that would be labeled by both strategies are present in the adult lung but not recognized in this data is not clear^32^. In a similar model of hyperoxia-induced neonatal lung injury, Cantu et al. noted a population of ‘reactive aCap’ EC that emerged in hyperoxia and shared Cap1 and Cap2 markers, likely representing Cap^INT^ EC^33^. Similarly, Vila Ellis et al. also identified a population of ‘early Cap2’ in a neonatal hyperoxia model, and a separate cluster of ‘transitional EC’ that emerged only in hyperoxia exposed mice lacking p53 in EC. In contrast to our results and those of Cantu, the ‘early’ Cap2 were not found to increase with hyperoxia, perhaps resulting from the much smaller number of EC sequenced in that dataset. Interestingly, although the ‘transitional’ EC shared some Cap1 and Cap2 markers, they lacked both *Plvap* and *Car4*, also in contrast to our data. Taken together, these data suggest that distinct versions of Cap^INT^ may emerge in different disease states, with unknown implications. Future studies will be important to better define how different injuries separately alter capillary fate and how distinct alterations to the capillary transcriptome induces discrete changes to the alveolar capillary niche.

Our study has several limitations, and important questions remain unanswered. First, the absence of unique markers defining the Cap^INT^ cell state limited our ability to perform genetic lineage tracing of these cells using easily accessible tools. Although RNA velocity suggested that Cap1 are the progenitors for both Cap^INT^ and Cap2, rather than Cap^INT^ serving as the progenitors for both Cap1 and Cap2, development of a dual, Cre-Dre reporter mouse model to permit lineage tracing of the Cap^INT^ will be essential for definitively answering this specific question. The long term-effects of disrupting Cap^INT^ to terminally differentiated Cap2 were not evaluated in the present report, including a lack of assessment for the development of pulmonary hypertension, definitive measures of lung fibrosis, or alterations in pulmonary function. Whether all Cap1 EC, or only a subset, have the capacity to serve as progenitors that can enter the intermediate state and then differentiate into Cap2 also remains unknown. Finally, although the physiologic adaptations accompanying birth appear to be a key driver of Cap2 differentiation, the molecular mechanisms that push Cap1 progenitors into the intermediate state were not identified in this study.

In summary, our study shows for the first time that differentiation of Cap1 progenitors into Cap2 EC in both development and disease requires transition through an intermediate cell state. Epithelial-derived VEGF has been previously shown to be required for Cap2 differentiation, however, in our study we show that VEGFR2 signaling is not required for entry into the intermediate cell state but is essential for transition from the Cap^INT^ to fully differentiated Cap2. Moreover, we show that Cap2 terminal differentiation continues after birth, and abrogating VEGFR2 signaling in Tbx2-expressing cells prevents Cap2 differentiation and increases cells that remain ‘trapped’ in the intermediate state. This abrogation both impairs alveolarization and appears to induce activation of alveolar fibroblasts to a myofibroblast-like phenotype, identifying a key role for terminally differentiated Cap2 in maintaining a homeostatic alveolar niche. Given that both postnatal alveolarization and lung regeneration require a coordinated balance of Cap1 and Cap2 growth, understanding the molecular mechanisms that regulated capillary EC plasticity has important implications from lung diseases characterized by alterations of capillary growth or disruption of the alveolar capillary niche.

## Methods

### Mice and experimental models

Wild-type C57BL/6 mice (Charles River Laboratories) were used as for developmental studies on wild-type mice. To generate mice permitting inducible deletion of VEGFR2 in Cap2 EC, Tbx2^tm1(EGFP/cre/ERT2)Wtsi^ mice (Tbx2-iCre; MGI:5633849) were purchased from the Jackson Laboratory and Vegfr2^flox/flox^ mice purchased from the Jackson Laboratory (JAX: 018977)^1,34^. For genetic lineage tracing, Tbx2-iCre mice were bred to Rosa26-tdTomato (Gt(ROSA)26Sor^tm14(CAG-tdTomato)Hze^)^35^ mice purchased from the Jackson Laboratory (JAX:007914). The day of birth was considered postnatal day 0 (P0). To induce Cre recombinase activity, tamoxifen (Sigma, T5648) was dissolved in corn oil and administered by intragastric injection (50μg/dose) once daily for three consecutive days (P0–P2) as previous described^36^. Neonatal mice were euthanized, and the lungs were perfused through the right ventricle with cold phosphate-buffered saline (PBS). Lungs were then pressure inflated with 4% paraformaldehyde (PFA) to preserve tissue architecture. After inflation, the lungs were harvested and fixed in 4% PFA at 4°C for 24 hours, as previously described as previously described^37^.

For studies employing chronic hyperoxia, newborn pups at P0 were exposed to 80% O₂ (hyperoxia) in a BioSpherix chamber (BioSpherix, Parish, NY) for 3, 7, or 14 days, to mimic the impaired alveolarization characteristic of BPD^38^. Dams were rotated every 24 hours. Lung cells or tissue were collected as described previously to assess hyperoxia effects.

### Fluorescence-activated cell sorting (FACS) of single cells

Mice were anesthetized and perfused as previously described. Lungs were immediately harvested and minced using scissors in a 60 mm Petri dish containing cold DMEM supplemented with 0.2 mg/ml Liberase (Roche) and DNase I. The tissue suspension was transferred into 15 ml tubes and incubated at 37°C for 15 minutes with shaking to facilitate enzymatic digestion. Digestion was halted by adding DMEM containing 10% fetal bovine serum (FBS), and the cell suspension was filtered through a 70 µm strainer to remove debris.

Cells were centrifuged at 500 × g for 5 minutes at 4°C and resuspended in red blood cell lysis buffer (Invitrogen eBioscience) for 3 minutes to remove erythrocytes. After lysis, cells were washed with 10 ml PBS and recentrifuged. The cell pellet was then resuspended and incubated with 10% FBS for 30 minutes on ice to block non-specific binding. For surface staining, cells were incubated for 30 minutes on ice with antibodies against CD31 (1:100, clone 390, Biolegend) and CD45 (1:100, clone F11, Biolegend), along with DAPI for viability assessment. The primary antibodies for cells were Car4 (1:100, AF2414, R&D), Kdr (1:100, AF644, R&D) and Kit (1:100, AF1356, R&D). Following staining, cells were washed twice with ice-cold PBS. If applicable, cells were incubated with the appropriate secondary antibody for 30 mins on ice. Endothelial cells were identified as DAPI⁻CD45⁻CD31⁺GFP⁺ or GFP⁻ populations and were sorted by FACS. The sorted endothelial cells (ECs) were subsequently processed for ATAC-seq. Flow data were analyzed using FlowJo software.

### Western immunoblot analysis

Whole lungs were dissected and tissue lysed in RIPA buffer (ThermoFisher, 89900) containing 1 mM PMSF and a phosphatase inhibitor cocktail (ThermoFisher). Samples were kept on ice for 10 minutes, then centrifuged at 13,000 rpm for 10 minutes at 4°C. The supernatant was collected, and protein concentration was measured using the Pierce BCA Protein Assay Kit (ThermoScientific, 23225) following the manufacturer’s protocol. Proteins were separated by electrophoresis and transferred onto a PVDF membrane. The membrane was blocked with 5% BSA (Sigma, A3059) in Tris-buffered saline for 1 hour at room temperature with agitation. Primary antibodies, α-SMA (Abcam, ab7817, 1:2000) or β-actin (Invitrogen, AM4302, 1:2000), were added and incubated overnight at 4°C. The membrane was washed three times with TBST, then incubated with an HRP-linked secondary antibody (1:2000) diluted in 2% BSA in TBST for 1 hour at room temperature with agitation. After washing, the membrane was treated with SuperSignal West Pico substrate (Thermo, 34580) for 1 minute and imaged using a BioRad ChemiDoc MP system.

### Immunofluorescence staining

Tissue sections or slides were fixed in 4% formalin (Sigma-Aldrich, St. Louis, MA, USA) and permeabilized with 1% Triton X-100 (Sigma-Aldrich). After blocking with 5% BSA for 1 hour, slides were incubated with the following primary antibodies: goat anti-CD31 (AF3628, R&D, 1:500), rabbit anti-AQP5 (ab78486, Abcam, 1:200), mouse anti-α-SMA Cy3 (C6198, Sigma-Aldrich, 1:200), rabbit anti-ERG (MA532036, Thermo Fisher Scientific,1:500). After primary incubation, slides were treated with fluorescence-conjugated secondary antibodies (Alexa Fluor 488, 568, or 647, Thermo Fisher, 1:500) and counterstained with DAPI (MBD0015, Thermo Fisher, 1:1000) to label nuclei. Controls were performed by omitting the primary antibodies.

### Real-time PCR

Total mRNA was isolated from sorted VEGFR^High^ EC from VEGFR2^WT^ mice and VEGFR2^Low^ EC from VEGFR2^ΔCap2^ mice using the RNeasy Micro Kit or Minikit (Qiagen, Valencia, CA, USA), and RNA concentration and purity assessed using a NanoDrop spectrophotometer. cDNA was synthesized using the HiScript II qRT SuperMix II (Vazyme, China). RT-PCR was performed with the 2X RT-PCR enzyme mix (Vazyme, China) and analyzed on a LightCycler 96 (Roche, Germany). RT-PCR primers used in this study were Taq-man primers Gapdh (thermofisher Mm99999915_g1), Kdr (Mm00440086_m1) and Vegfr1 (Mm00438980_m1).

### Single-nucleus RNA-sequencing

Lung tissue from four experimental groups Kdr^flox/flox^ and Tbx2^icre^Kdr^flox/flox^ under maintained in normoxia or exposed to chronic hyperoxia were euthanized at P7. Lung tissue was dissected and snap-frozen in liquid nitrogen. Nuclei were prepared from frozen tissue using a nuclear isolation kit from 10X Genomics (10x Genomics, PN-1000494). For 10X Chromium, 10,000 nuclei were loaded per lane. The libraries were prepared according to manufacturer protocol (10X Genomics) using 3’V3 Kits (10X Genomics, PN-1000269) and were submitted for sequencing via a NovaSeq 6000 flow cell to a depth of 50,000 reads/cell. Raw sequencing data were processed using CellRanger pipeline.

### Single cell/single nucleus sequencing data analysis

Sequencing reads were aligned to the Grcm39 mouse genome using Cellranger (v9.0.0). SoupX was used to remove ambient RNA signal^39^. Gene expression count tables were turned into anndata objects and processed using scanpy when not otherwise specified^40^. Cells that had more than five median absolute deviations of either unique genes or UMIs detected were removed. Cells that had more than three median absolute deviations of percent mitochondrial or ribosomal UMIs were removed^41^. Scrublet was used to automatically detect and remove doublets^42^. Counts were normalized and log transformed. PCA was run prior to embedding and clustering. The Leiden algorithm was used for clustering, and Uniform Manifold Approximation and Projection for embedding^43^. Lineage identity was assigned using canonical markers: *Cdh5* (endothelial); *Epcam* (epithelial); *Ptprc* (immune); and *Col1a1* (mesenchymal). Cell typing was performed after each lineage was re-clustered and embedded as detailed above. Cell types were assigned a second time after regressing out cell cycle genes. RNA velocity was determined using velocyto to align reads, and scVelo to calculate trajectories^44,45^. For more detail see https://github.com/CarstenKnutsen/capillary_differentiaion_manuscript

### Assay for transposase-accessible chromatin using sequencing (ATAC-seq) and data analysis

ATAC-seq was performed using the Active Motif ATAC-seq Kit (Cat# 53150) according to the manufacturer’s instructions. Approximately 50,000 EC were processed per sample. Cells were washed with 100 µl of ice-cold PBS and centrifuged at 500 × g for 5 minutes at 4°C. The cell pellet was resuspended in 100 µl of pre-chilled ATAC lysis buffer and immediately centrifuged again at 500 × g for 10 minutes at 4°C. The resulting nuclei pellet was resuspended in the Tagmentation Master Mix and incubated at 37°C for 30 minutes in a thermomixer. Following tagmentation, DNA was purified according to the protocol. Library amplification was performed as recommended and purified using SPRI bead cleanup. Sequencing was conducted using paired-end 150 bp reads. Sequencing reads were mapped using the nfcore/atacseq pipeline of the mouse reference genome^46^ (mm39). Peak calling was performed using MACS2, and differential chromatin accessibility analysis was carried out using HOMER and DESeq2.

### Multiplex fluorescent in situ hybridization

Post-natal mice were euthanized as described earlier. Lung tissue was placed in 10% neutral buffered formalin and incubated at 4°C for 20 hours. After fixation, lungs were washed twice in 1x PBS and transferred to 70% ethanol for paraffin embedding. The RNAscope Multiplex Fluorescent v2 Assay kit (Advanced Cell Diagnostics) was used according to the manufacturer’s instructions. Formalin-fixed paraffin-embedded (FFPE) lung sections (5 μm) were used within a day of sectioning for optimal results. Nuclei were counterstained with DAPI (Life Technology Corp.). Signal amplification was achieved using Opal dyes (Akoya Biosciences) via the manufacturer’s protocol. Probes (Advanced Cell Diagnostics) used were Mm-Tbx2 (448991), Mm-Kit-C2 (314151-C2), Mm-Plvap-C3 (440221-C3). Images were captured using Leica SP8 confocal microscope with 405nm, 488nm, 560nm and 633nm excitation lasers. Both merged and split channel images were collected.

For quantification of Cap^INT^ abundance, Cap^INT^ cells were identified using Tbx2, Kit, and Plvap probes. Cells with three or more puncta were considered positive. More than three random fields of view were imaged, and over 400 nuclei were counted. Each group had at least two replicates.

### Statistical Analysis

All data are presented as mean ± SEM. Statistical differences between two groups were determined using Student’s t-test. A P-value ≤ 0.05 was considered statistically significant.

## Acknowledgements

This study was supported by funding from the Marc and Lynne Benioff Endowed Chair (C.M.A.) and NIH NHLBI R01-HL154002, HL155828, and HL160018; C.M.A.). We would like to acknowledgement support from the Stanford Shared FACS facility and Stanford Cell Sciences Imaging Facility, with particular thanks to Kitty Lee. The results here in Fig. 4E are based upon data generated by the LungMAP Consortium and downloaded from (www.lungmap.net), in May, 2025. The LungMAP consortium, the Human Tissue Core (U01-HL144861), and the LungMAP Data Coordinating Center (U24-HL148865) are funded by the National Heart, Lung, and Blood Institute (NHLBI).

## Data Availability

Single cell RNA sequencing data generated for this dataset can be found at https://figshare.com/articles/dataset/P7_Kdr-KO_scRNA_seq_data/29626994?file=56490755. ATAC sequencing data can be found at https://figshare.com/articles/dataset/P7_Cap2_and_EC_ATAC_data/29627333. Early postnatal data (P1-P21) was sourced from https://figshare.com/articles/dataset/Cell_atlas_of_the_murine_perinatal_lung/14703792?file= 28238682. Multiple datasets from the Gene Expression Omnibus (GEO) were used including E18 and P0 ECs from GSE149563, and P3-P14 ECs under hyperoxia from GSE151974. P3 EC data was sourced from https://figshare.com/account/articles/29079218?file=54578297. Human neonatal data was sourced from https://cellxgene.cziscience.com/collections/28e9d721-6816-48a2-8d0b-43bf0b0c0ebc. Human BPD data was sourced from LungMap dataset code LMEX0000004400 at https://www.lungmap.net/dataset/?experiment_id=LMEX0000004400.

---

**Figure S1:**
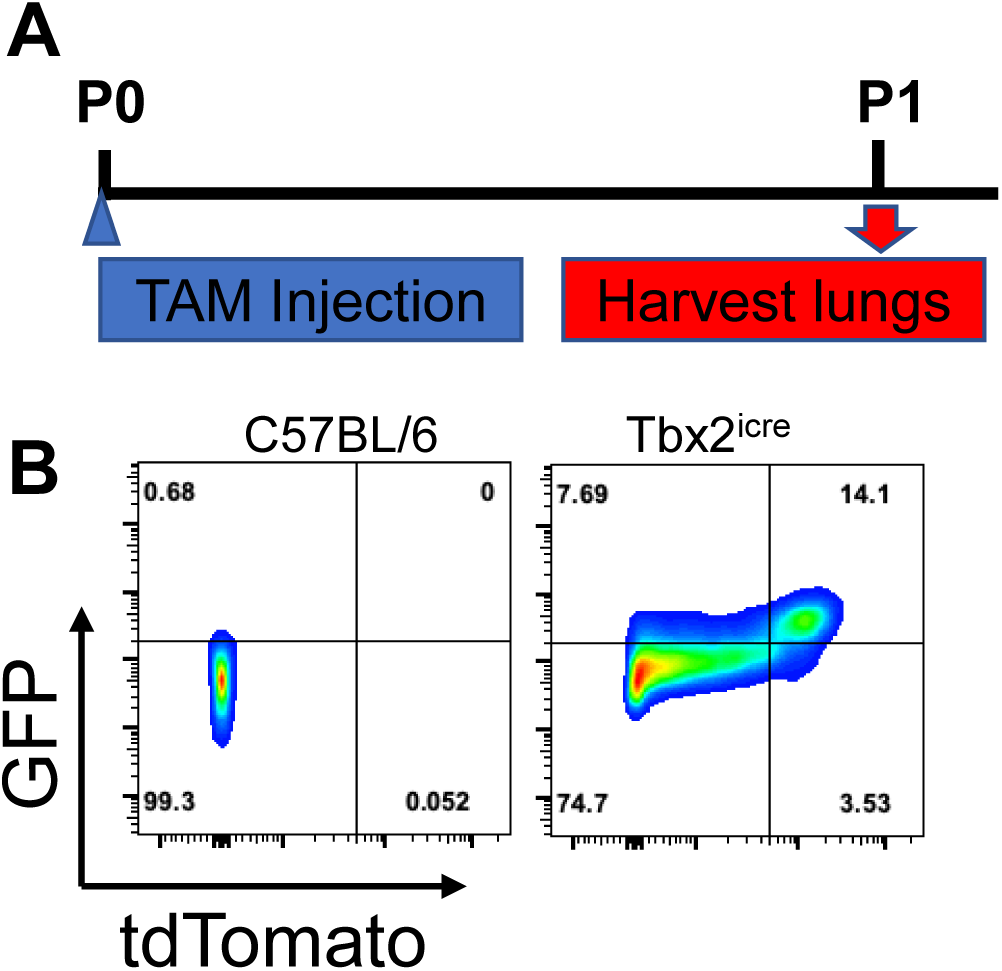
Efficiency of tdT labeling by Tbx2^icre^ in Tbx2-expressing ECs. (A) Schematic of the tamoxifen administration protocol in the Tbx2-iCre-tdTomato mice. (B) Representative FACS plot showing the separation of CD31+ EC by levels of tdT (x-axis) or GFP (y-axis) expression.

**Figure S2:**
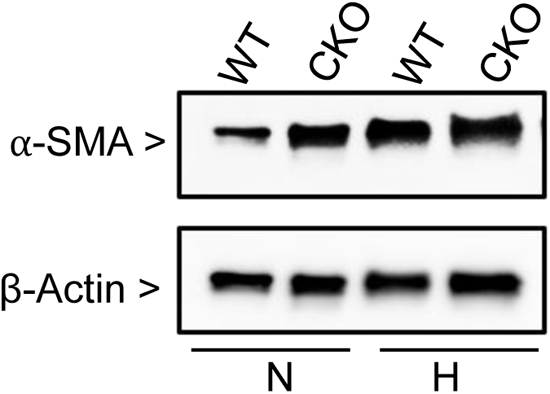
Postnatal loss of VEGFR2 from Tbx2+ EC increases a-SMA expression. WB from whole lung tissue obtained from lung tissue from VEGFR2^WT^ and VEGFR2^DCap2^ mice at P7 exposed to normoxia (N) or hyperoxia (H).

**Figure S3:**
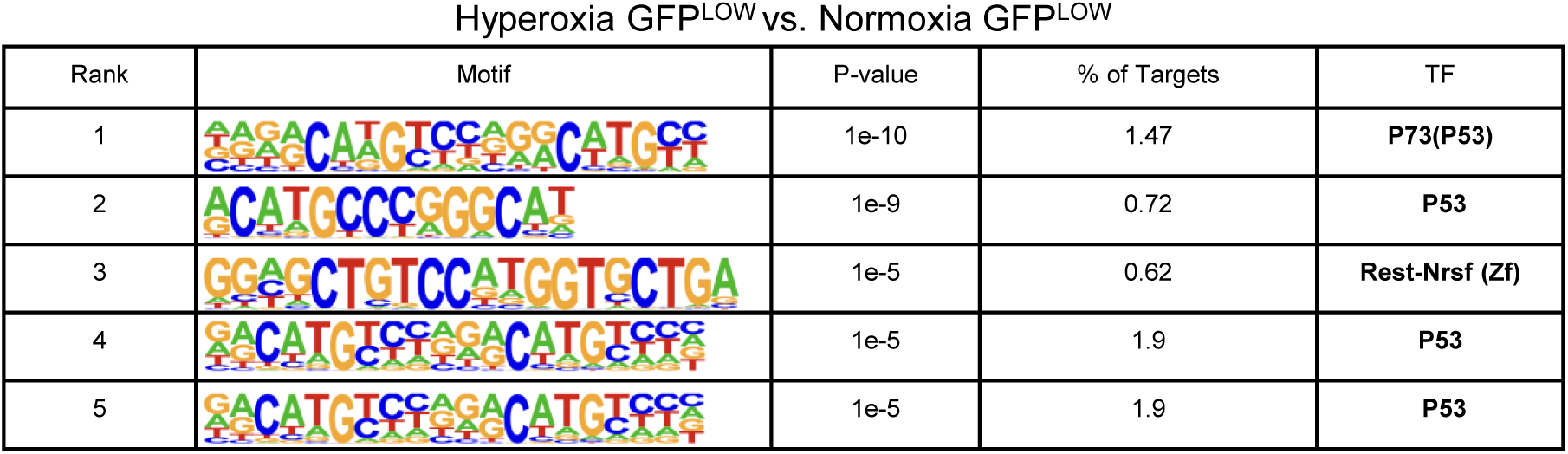
Table including the top predicted enriched TF binding domains in the DAPs in the hyperoxic versus normoxic GFP^LOW^ EC.

## Notes

### Competing Interest Statement

The authors have declared no competing interest.

### Summary of Updates

Author affiliations updated; Figures 4,7 updated.

## References

1. Bhatt, A. J. et al. Disrupted Pulmonary Vasculature and Decreased Vascular Endothelial Growth Factor, Flt-1, and TIE-2 in Human Infants Dying with Bronchopulmonary Dysplasia. Am. J. Respir. Crit. Care Med. 164, 1971–1980 (2001).

2. Jakkula, M. et al. Inhibition of angiogenesis decreases alveolarization in the developing rat lung. Am. J. Physiol.-Lung Cell. Mol. Physiol. 279, L600–L607 (2000).

3. Thébaud, B. et al. Vascular Endothelial Growth Factor Gene Therapy Increases Survival, Promotes Lung Angiogenesis, and Prevents Alveolar Damage in Hyperoxia-Induced Lung Injury: Evidence That Angiogenesis Participates in Alveolarization. Circulation 112, 2477– 2486 (2005).

4. Toya, S. P. & Malik, A. B. Role of endothelial injury in disease mechanisms and contribution of progenitor cells in mediating endothelial repair. Immunobiology 217, 569–580 (2012).

5. Thane, K., Ingenito, E. P. & Hoffman, A. M. Lung regeneration and translational implications of the postpneumonectomy model. Transl. Res. 163, 363–376 (2014).

6. Jancelewicz, T., Grethel, E. J., Chapin, C. J., Clifton, M. S. & Nobuhara, K. K. Vascular Endothelial Growth Factor Isoform and Receptor Expression During Compensatory Lung Growth. J. Surg. Res. 160, 107–113 (2010).

7. Rafii, S., Butler, J. M. & Ding, B.-S. Angiocrine functions of organ-specific endothelial cells. Nature 529, 316–325 (2016).

8. Ding, B.-S. et al. Endothelial-Derived Angiocrine Signals Induce and Sustain Regenerative Lung Alveolarization. Cell 147, 539–553 (2011).

9. Ding, B.-S. et al. Inductive angiocrine signals from sinusoidal endothelium are required for liver regeneration. Nature 468, 310–315 (2010).

10. Hooper, A. T. et al. Engraftment and Reconstitution of Hematopoiesis Is Dependent on VEGFR2-Mediated Regeneration of Sinusoidal Endothelial Cells. Cell Stem Cell 4, 263– 274 (2009).

11. Ding, B.-S. et al. Divergent angiocrine signals from vascular niche balance liver regeneration and fibrosis. Nature 505, 97–102 (2014).

12. Jiang, H. et al. Angiocrine FSTL1 (Follistatin-Like Protein 1) Insufficiency Leads to Atrial and Venous Wall Fibrosis via SMAD3 Activation. Arterioscler. Thromb. Vasc. Biol. 40, 958–972 (2020).

13. Gillich, A. et al. Capillary cell-type specialization in the alveolus. Nature 586, 785–789 (2020).

14. Vila Ellis, L., et al. Epithelial Vegfa Specifies a Distinct Endothelial Population in the Mouse Lung. Dev. Cell 52, 617–630.e6 (2020).

15. Zanini, F. et al. Developmental diversity and unique sensitivity to injury of lung endothelial subtypes during postnatal growth. iScience 26, 106097 (2023).

16. Hurskainen, M. et al. Single cell transcriptomic analysis of murine lung development on hyperoxia-induced damage. Nat. Commun. 12, 1565 (2021).

17. Negretti, N. M. et al. A single-cell atlas of mouse lung development. Development 148, dev199512 (2021).

18. Niethamer, T. K. et al. Defining the role of pulmonary endothelial cell heterogeneity in the response to acute lung injury. eLife 9, e53072 (2020).

19. Ellis, L. V., Bywaters, J. D. & Chen, J. Endothelial deletion of p53 generates transitional endothelial cells and improves lung development during neonatal hyperoxia. bioRxiv (2024).

20. Sveiven, S. N., Knutsen, C., Zanini, F., Cornfield, D. & Alvira, C. Single Cell RNA Sequencing Reveals Gene Expression Continuums Along the Spatial Hierarchy of the Pulmonary Circulation. bioRxiv (2025).

21. Bhattacharya, S. et al. Single-Cell Transcriptomic Profiling Identifies Molecular Phenotypes of Newborn Human Lung Cells. Genes 15, 298 (2024).

22. Auyeung, V. C. et al. IRE1α drives lung epithelial progenitor dysfunction to establish a niche for pulmonary fibrosis. Am. J. Physiol.-Lung Cell. Mol. Physiol. 322, L564–L580 (2022).

23. Choi, J. et al. Inflammatory Signals Induce AT2 Cell-Derived Damage-Associated Transient Progenitors that Mediate Alveolar Regeneration. Cell Stem Cell 27, 366–382.e7 (2020).

24. Zou, M. et al. Latent Transforming Growth Factor-β Binding Protein-2 Regulates Lung Fibroblast-to-Myofibroblast Differentiation in Pulmonary Fibrosis via NF-κB Signaling. Front. Pharmacol. 12, 788714 (2021).

25. Kurundkar, A. R. et al. The matricellular protein CCN1 enhances TGF-β1/SMAD3-dependent profibrotic signaling in fibroblasts and contributes to fibrogenic responses to lung injury. FASEB J. 30, 2135–2150 (2016).

26. Raza, S. et al. SOX9 is required for kidney fibrosis and activates NAV3 to drive renal myofibroblast function. Sci. Signal. 14, eabb4282 (2021).

27. Thickett, D. R., Armstrong, L. & Millar, A. B. A Role for Vascular Endothelial Growth Factor in Acute and Resolving Lung Injury. Am. J. Respir. Crit. Care Med. 166, 1332–1337 (2002).

28. Kanazawa, H., Asai, K., Hirata, K. & Yoshikawa, J. Possible effects of vascular endothelial growth factor in the pathogenesis of chronic obstructive pulmonary disease. Am. J. Med. 114, 354–358 (2003).

29. Shah, A. V. et al. The endothelial transcription factor ERG mediates Angiopoietin-1-dependent control of Notch signalling and vascular stability. Nat. Commun. 8, 16002 (2017).

30. Desai, T. J., Brownfield, D. G. & Krasnow, M. A. Alveolar progenitor and stem cells in lung development, renewal and cancer. Nature 507, 190–194 (2014).

31. Choi, J. et al. Inflammatory Signals Induce AT2 Cell-Derived Damage-Associated Transient Progenitors that Mediate Alveolar Regeneration. Cell Stem Cell 27, 366–382.e7 (2020).

32. Niethamer, T. K. et al. Longitudinal single-cell profiles of lung regeneration after viral infection reveal persistent injury-associated cell states. Cell Stem Cell 32, 302–321.e6 (2025).

33. Cantu, A. et al. Remarkable sex-specific differences at Single-Cell Resolution in Neonatal Hyperoxic Lung Injury. bioRxiv.

34. Hooper, A. T. et al. Engraftment and Reconstitution of Hematopoiesis Is Dependent on VEGFR2-Mediated Regeneration of Sinusoidal Endothelial Cells. Cell Stem Cell 4, 263– 274 (2009).

35. Madisen, L. et al. A robust and high-throughput Cre reporting and characterization system for the whole mouse brain. Nat. Neurosci. 13, 133–140 (2010).

36. Rao, S., Liu, M., Iosef, C., Knutsen, C. & Alvira, C. M. Endothelial-specific loss of IKKβ disrupts pulmonary endothelial angiogenesis and impairs postnatal lung growth. Am. J. Physiol.-Lung Cell. Mol. Physiol. 325, L299–L313 (2023).

37. Domingo-Gonzalez, R. et al. Diverse homeostatic and immunomodulatory roles of immune cells in the developing mouse lung at single cell resolution. eLife 9, e56890 (2020).

38. Hilgendorff, A., Reiss, I., Ehrhardt, H., Eickelberg, O. & Alvira, C. M. Chronic Lung Disease in the Preterm Infant. Lessons Learned from Animal Models. Am. J. Respir. Cell Mol. Biol. 50, 233–245 (2014).

39. Young, M. D. & Behjati, S. SoupX removes ambient RNA contamination from droplet-based single-cell RNA sequencing data. GigaScience 9, (2020).

40. Wolf, F. A., Angerer, P. & Theis, F. J. SCANPY: large-scale single-cell gene expression data analysis. Genome Biol. 19, (2018).

41. Heumos, L. et al. Best practices for single-cell analysis across modalities. Nat. Rev. Genet. 24, 550–572 (2023).

42. Wolock, S. L., Lopez, R. & Klein, A. M. Scrublet: Computational Identification of Cell Doublets in Single-Cell Transcriptomic Data. Cell Syst. 8, 281–291.e9 (2019).

43. Traag, V. A., Waltman, L. & Van Eck, N. J. From Louvain to Leiden: guaranteeing well-connected communities. Sci. Rep. 9, (2019).

44. La Manno, G. et al. RNA velocity of single cells. Nature 560, 494–498 (2018).

45. Bergen, V., Lange, M., Peidli, S., Wolf, F. A. & Theis, F. J. Generalizing RNA velocity to transient cell states through dynamical modeling. Nat. Biotechnol. 38, 1408–1414 (2020).

46. Ewels, P. A. et al. The nf-core framework for community-curated bioinformatics pipelines. Nat. Biotechnol. 38, 276–278 (2020).

